# Climate resilience conserved in global germplasm repositories: Picking the most promising parents for agile plant breeding

**DOI:** 10.1101/2024.05.11.593573

**Authors:** Quinn Campbell, Nora Castaneda-Alvarez, Ryan Domingo, Eric Bishop-von Wettberg, Bryan Runck, Hervé Nandkangré, Anna McCormick, Nathan Fumia, Jeffrey Neyhart, Benjamin Kilian, Peterson Wambugu, Desterio Nyamongo, Sariel Hübner, Sidney Sitar, Addie Thompson, Loren Rieseberg, Michael A. Gore, Michael Kantar

**Affiliations:** Department of Tropical Plant and Soil Sciences, University of Hawai’i at Mānoa, Honolulu, HI, USA; Global Crop Diversity Trust, Bonn, Germany; Department of Plant and Soil Science, University of Vermont, Burlington, VT, USA; Department of Geography, Environment, and Society, University of Minnesota, Minneapolis, MN, USA; USDA-ARS, Genetic Improvement for Fruits and Vegetables Lab, Chatsworth, NJ, USA; Kenya Agricultural and Livestock Research Organization | KALRO, Genetic Resources Research Institute, Kikuyu, Kenya; Galilee Research Institute (MIGAL), Tel Hai College, Upper Galilee, Israel; Department of Plant, Soil, and Microbial Sciences, Michigan State University, East Lansing, Michigan, USA; Department of Botany, University of British Columbia, Vancouver, Canada; School of Integrative Plant Science, Cornell University, Ithaca, New York, USA

**Keywords:** Genomic Selection, FIGS, Future Projection, Climate Resilience, Crop Adaptation

## Abstract

Crop diversity is an essential resource for national and international breeding programs aimed at preparing global agriculture for a changing climate to ensure global food security. To do this there are related risks that need to be evaluated (1) does the genetic diversity needed for climate adaptation exist somewhere? And (2) is such genetic diversity accessible? To evaluate these risks, we consider the test case of publicly available genotyped and georeferenced sorghum landraces (n = 1,937) to ask if diversity is sufficient to support breeding for climate change adaptation. Answering these questions allows for characterization of the best potential parents and the geographies that harbor the most potentially promising genetypes for crop improvement. We subset this data into national, regional, and global geographic regions, and complete/mini core collections to understand the potential for climate adaptation in regional germplasm. Study accessions were given a future climate resilience score based on future climatic projections and a genomic adaptive capacity score using genomic estimated adaptive values (GEAVs) generated from environmental genomic selection - EGS) to ask whether this accessible diversity stored in germplasm repositories is potentially sufficient to meet forecasted changes in growing environments under climate change. We find that genomic resilience capacity is highly variable among countries and regions. High geographical variability was also found for climate resilience. To equitably adapt agriculture to future climate conditions, increased accessibility to plant genetic resources is essential.

## Main

Plant genetic resources are foundational to food system climate adaptation [1-3]. Breeders leverage genetic diversity to adapt crops to a changing climate by identifying traits or lines from germplasm repositories, farmers’ fields, or other breeding programs that exhibit climate resilient variation [4-7]. However, collection utility is dependent on characterization, funding and access [8-13], and while plants, animals and diseases do not recognize national borders, phytosanitary regulations that are the mainstay of nation states to limit the spread of potentially infectious diseases for nearly 80 years [14], can severely limit breeders’ access to novel genetic variation for adapting local breeding programs to climate change. The Nagoya Protocol to the Convention on Biological Diversity and the International Treaty for Plant Genetic Resources for Food and Agriculture (ITPGRFA) has allowed countries to create bilateral arrangements for germplasm access and benefit sharing, which has facilitated access to genetic resources [15]. However, the ITPGRFA, which established a multilateral system for germplasm exchange, does not cover all major staple crops (e.g., soybean, sugar cane, oil palm and groundnut). Thus, for non-treaty crops, bilateral arrangements are required for access. However, there are ongoing negotiations at the Treaty’s Governing Body to review the list of species covered under the Treaty’s multilateral system and once complete, which could address this challenge. Despite this progress, phytosanitary restrictions, national/international regulations, and administrative complications remain major limiting factors in the use of genetic resources for breeding for abiotic stresses under a future climate [16]. These necessary hindrances to germplasm exchange create a need for breeders and policy-makers to have access to improved information on the potential performance of lines in their local environment.

### Climate change impact on cultivated plants

Achieving climate change adaptation and mitigation through plant breeding requires both that the genetic variation exists within a crop gene pool and that this germplasm is accessible to plant breeders. These two challenges - one biological and the other social/regulatory/operational - set the broad context required to evaluate the adaptation potential of different countries’ agricultural systems under climate change. This body of work explores a range of strategies associated with adapting food systems to climate change, both future biotic and abiotic threats [17-18]. Presently, academic and grey literature have focused primarily on the biological risk [19-21]. Thus the need to adapt crops to climate change is a major impetus for preventing crop genetic erosion [3]. Different methods have been proposed to identify the best accessions from collections for use in breeding - these include core collections [22-23], Focused Identification of Germplasm Strategy (FIGS) [24], landscape genomics [25-27], and germplasm genomics [28]. These strategies aim to reduce the number of accessions to be evaluated, and subsequently deployed in crosses, thereby increasing the efficiency of the pre-breeding process [29].

Here, we present an approach for considering the biological and social risks of crop adaptation at both global and national levels. Our approach combines germplasm genomics with geographic indices that can be used for national and sub-national decision-making. We apply this approach to the example case of sorghum, a staple crop for subsistence farmers in rain-fed systems across sub-Saharan Africa [30], which shows great potential for adaptation to novel environments.

### Environmental Genomic Selection using the mini core as the training population

The United States National Plant Germplasm System (NPGS) and the International Crops Research Institute for the Semi-Arid Tropics (ICRISAT) sorghum collections represent the two largest collections of sorghum globally, and each contain accessions sourced from all major sorghum growing areas (**Table S1 & S2**). These collections came from regions that have extensive environmental and genetic variation (**Figure S1 & S2**). From these two major collections, Lasky et al. (2015) [4] sequenced a single genotype from each of 1,943 accessions chosen to maximize geographic representation. Of these, 1,937 genotypes (hereafter, study panel) form the basis of the present study. To operationalize this variation among the 1,937 study genotypes for use in adaptation to climate change, it is important to provide ways to partition the collection for specific traits of interest that fit different goals. Here we characterize the value of specific genotypes as parents using an analog to the genomic estimated breeding value (GEBV) - the genomic estimated adaptive value (GEAV), where climate and soil information is substituted for observed traits to predict heritable environmental adaptation [31] We utilize the sorghum mini core collection developed by Upadhyaya et al. (2009) as the training population [32]. Because the study genotypes are samples of landrace accessions minimally regenerated through selfing, it is reasonable to assume that they were under selection for hundreds of generations to enhance adaptation to the local environment. We characterize the resilience potential of national, regional and global germplasm collections based on this assumption. Here the predicted genome-wide value of genotypes as parents for adaptation to specific bioclimatic and biophysical features was assessed. Overall, there was predictive ability greater than r =0.5 for most of the environmental variables (**Figure S3**). Evaluating the potential of germplasm for a local breeding program before navigating phytosanitary and other restrictions can be understood at the individual molecular marker (e.g. single nucleotide polymorphism - SNP), chromosomal, or population genomic level (genomic estimate of resilience -GEAV) (**Figure 1-2**). Here this is based on the summed marker effects (SNPs) over chromosomes with genotypes organized by country of origin (**Figure 1)**.

**Figure 1.**
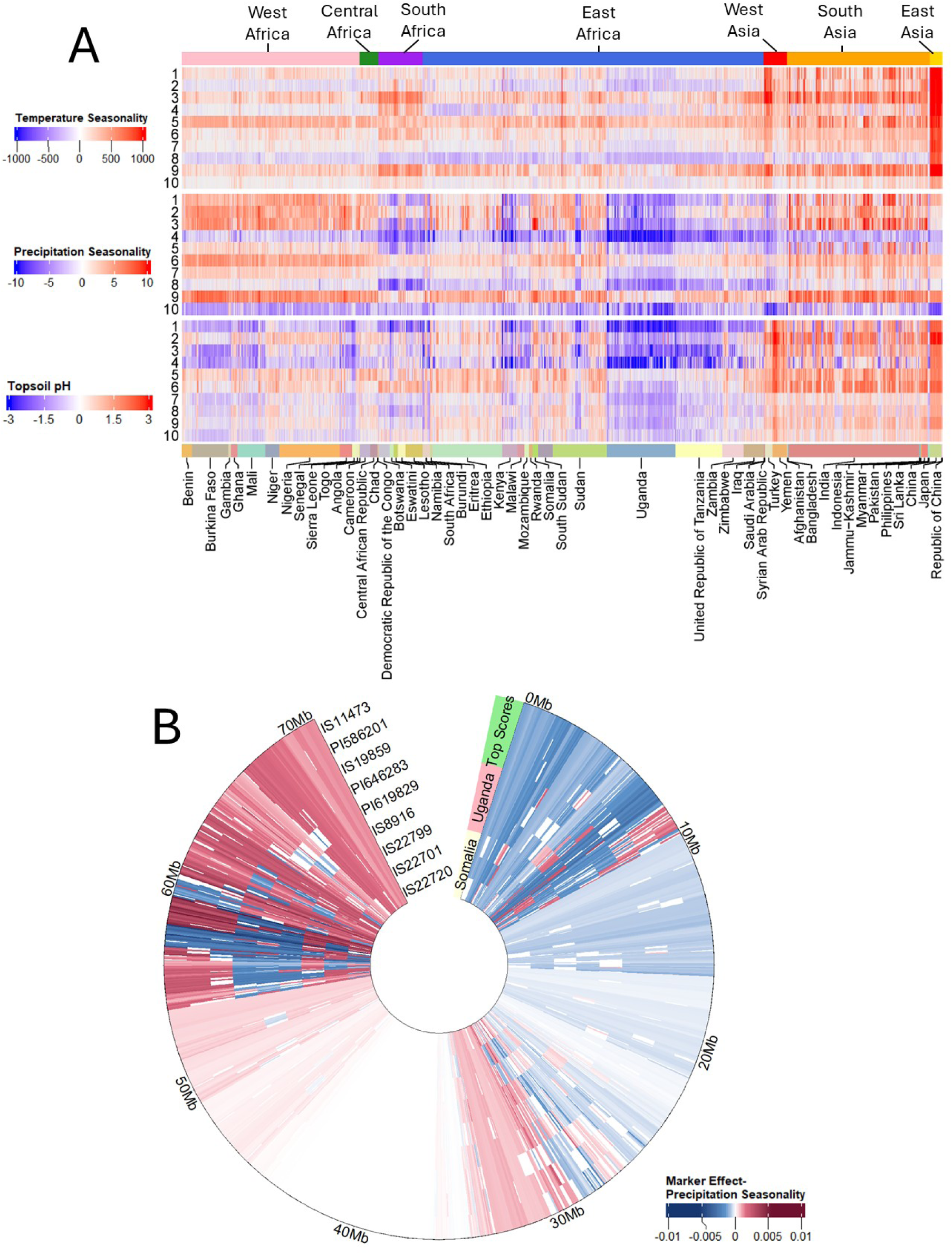
Here we present the genomic value of a specific chromosome for each line, specifically the predicted genetic value for a given environmental context. Marker effects for each chromosome were explored among genotypes originating from different geographic localities. A) Heatmap of chromosomal effects for the 10 sorghum chromosomes with environmental and soil variables. Columns represent the 1,937 sequenced sorghum genotypes, organized by country (below) and region (above) of original collection. Rows represent separate chromosomes. Values are the sum of marker effects (404,627 SNPs) from the genomic prediction across each chromosome. The three variables plotted here (temperature seasonality, precipitation seasonality, and topsoil pH) represent high genomic prediction accuracy from their respective category (**Figure S3**). B) Heatmap of marker effects on precipitation seasonality for 1,115 SNPs (filtered for LD of 0.20) on chromosome 3. Genotypes shown, from outside in, include: the three genotypes with the highest adaptive capacity, three genotypes from Uganda, and three genotypes from Somalia.

**Figure 2.**
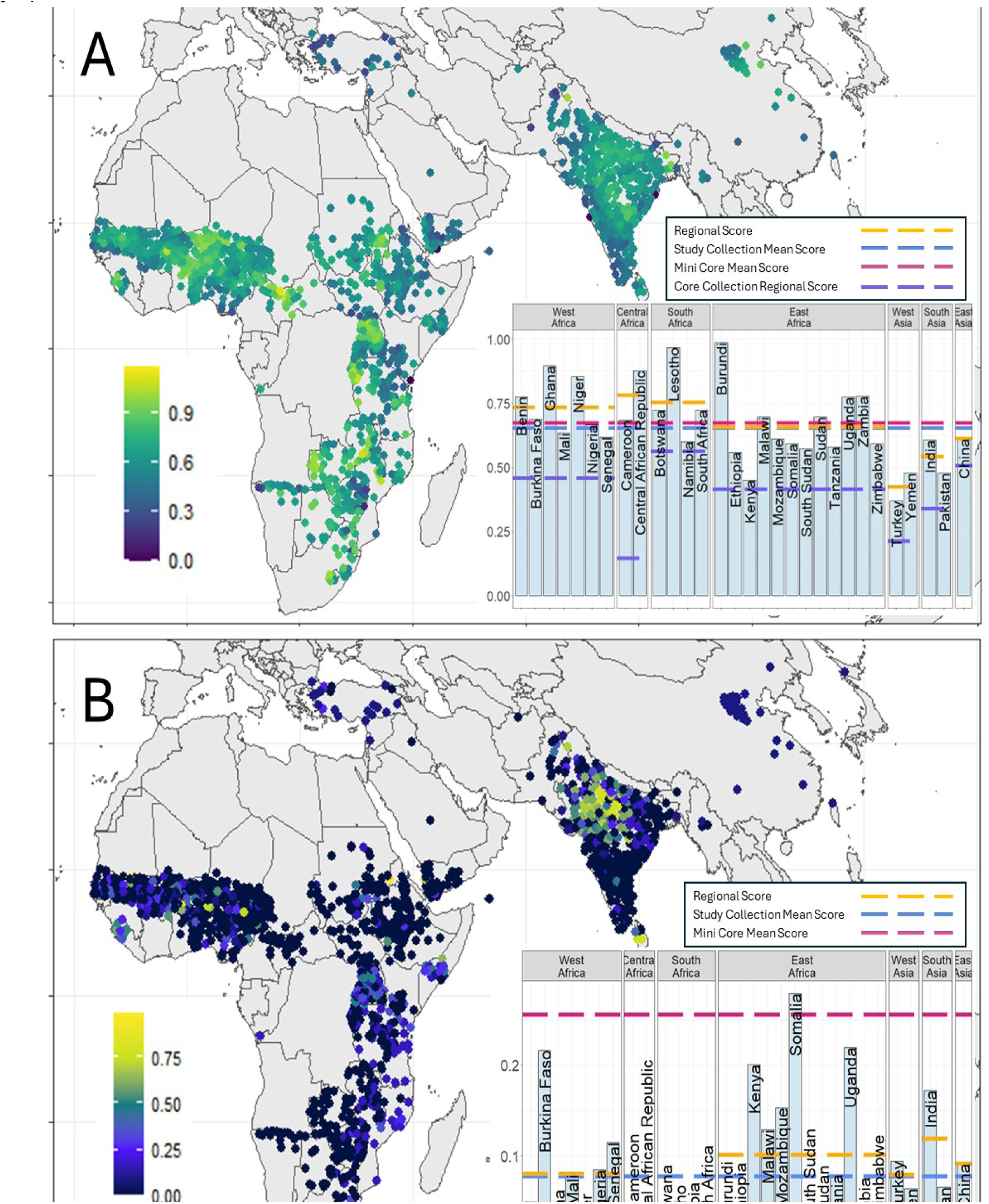
Barcharts of scores by country and region for A) Future Climate Resilience Score. Distribution of genotypes that are above future thresholds in important growing regions. Individual country scores with regional means (blue dotted line) and global median (red dotted line) showing areas with climate resilient genotypes and areas where genotypes are not resilient under climate change. B) Genomic Adaptive Capacity Score. Distribution of genotypes across provenances for Environmental Genomic Selection - EGS. Individual country scores with regional means (blue dotted line) and global median (red dotted line) showing the areas with broadly climate-adapted genotypes. Genotypes with higher scores can be thought of as generalists, they should be broadly adapted to many different potential climates. Genotypes with lower scores should be thought of as specialists, they will not be broadly adapted but may be useful in a very specific climate.

In this case, the chromosomes of certain genotypes are predicted to have higher GEAVs for specific abiotic/soil stressors (**Figure 1A**). For example, chromosome 8 is generally negatively associated with temperature seasonality, but this pattern is reversed for sorghum genotypes from China. These genotypes represent the northernmost point of origin among study genotypes, and therefore the environment with the highest temperature seasonality. Extending this analysis across the germplasm collection allows for identification of contrasting GEAVs, and thus promising individuals, geographic areas, and genomic regions for both sources and targets of crop improvement. Further, in genotypes from Tanzania and Uganda, the entire genome tends to be associated with lower precipitation seasonality and lower pH soils, while in other geographic regions patterns are more complex (**Figure 1A**). These patterns of contrasting GEAVs are also evident across chromosomes when looking at individual marker effects: for example, on chromosome 3 the genotypes with the top-ranking GEAVs [32] across climatic variables show distinctly different patterns of marker effects for precipitation seasonality when compared to genotypes from Somalia, which have positive chromosomal effects at chromosome 3, and genotypes from Uganda, which have negative summed marker effects at chromosome 3 (**Figure 1B**). Examples can be found for nearly every bioclimatic/biophysical trait examined. Thus, this approach that relies on a well-known statistical technique and easily obtained data identifies the most promising genotypes to be used as parents for any of these potential abiotic stressors.

### Climate Resilience and Adaptive Capacity Scores based on Geolocation and Genomics

The future climate resilience score for a particular genotype shows the proportion of cropland under sorghum cultivation within the country of origin where the present climate at the genotype’s collection site will be present in 2050, using CMIP6 models under the SSP 585 scenario. SSP Scenario 585 considers growing integration across global markets and heavy reliance on fossil fuels. National and regional scores are the mean scores of all genotypes collected in the given geographical area. Using this empirical outlier approach, we explored climate resilience partitioned by global, regional, and national provenance (**Figure 2a**). The score given to a region or country represents the maximum extent of present cropland conditions to which existing genotypes will remain suited in the future. The overall interpretation is that countries or regions with high scores will have less of a need to import germplasm for climate change adaptation compared to countries with the low scores. Analyzing the study genotypes in this way allows for the exploration of which genotypes may have the potentially promising use in breeding for specific abiotic stressors that will be most relevant under future projected climates. This method does not consider genetic information, only environmental data for the location where the accession that the genotype was sampled from was originally collected. For the study genotypes, global mean resilience was 0.746, regional resiliency ranged from 0.422-0.781, and national resilience ranged from 0.368-0.987. Central Africa had the highest regional climate resilience, while West Africa and South Africa were close behind with Burundi having the highest national resilience. There was not a meaningful difference in resilience between the mini core collection (mean = 0.731), the full core collection (determined by Grenier et al, 2001 [33]; hereafter, core collection), (mean = 0.724) and the full study panel (mean = 0.746). This score may also indicate that climate within the country is not predicted to change significantly, so current landraces will still be within the range of abiotic stress tolerance in the coming decades. However, this does not change our interpretation of the score, as genotypes from regions not predicted to experience large climatic changes still can be said to be resilient under climate change, while low scores identify genotypes with low climate resiliency and regions lacking well-adapted germplasm under future climates.

In our analysis, we consider two scenarios of germplasm availability: one in which germplasm is readily available globally, and a second one with restricted exchange of germplasm that assumes that germplasm is only available within its country of origin. The climate resilience score suggests that some genotypes will still be useful for climate adaptation in 2050. While there will obviously be maladaptation for a small proportion of the genotypes, particularly in the case of novel climates, there is potentially enough diversity to help adapt cultivated types. In a scenario of restricted germplasm exchange the spatial variation in climatic conditions within countries leads to broad adaptability amongst the genotypes implying that even where climatic conditions change as it is predicted/expected to, it is still possible to obtain genetic materials with the necessary adaptive capacity from within the country. In case of novel climates, genotypes available in the overlap areas within a country will harbor potentially useful variation to adapt in the future climates.

Further, we extend the use of EGS to create a genomic adaptive capacity score to explore the value of different genotypes (**Figure 2b**). This score provides multiple types of information for breeders. First, for those geographies that score high, it suggests to breeders where there are genotypes with potentially broadscale adaptation. Second, for those areas with a moderate score, it suggests that germplasm may be regionally adapted, and thus be useful to neighboring countries. Lastly, the locations scoring lowest predict the highest levels of local adaptation.

Using these general interpretive benchmarks, we observe that global adaptive capacity was 0.103, whereas regional resiliency ranging from 0.003-0.118 and national adaptive capacity ranging from 0-0.279. The region with the lowest genomic adaptive capacity was South Africa (0.166). The country with the highest genomic adaptive capacity was Somalia (0.279). Further, we see a large concentration of genotypes with high genomic adaptive capacity around the Punjab region of India, indicating the large capacity for broad climate resilience for genotypes collected from this region. This genomic adaptive capacity score showed a different pattern than the future climate resilience score (**Figure S6**), indicating that bioclimatic/biophysical and genetic data provide different information regarding which geographies (regional and national) may provide the best parental material for breeding in response to climate change.

### Using the Indices for Decision Support

Burundi had the highest national future climate resilience score (the maximum extent of present cropland conditions to which existing genotypes will remain suited in the future) based on study genotypes collected in the country; however, it also showed the lowest genomic adaptive capacity score based on genetic data (lowest proportion of genotypes that can serve as promising parents). This implies that the country is potentially well-positioned to respond to climate change using germplasm collected within the country. However, the low genomic adaptive capacity score means that genotypes in the study panel that originated in Burundi may not be as valuable internationally for breeding in response to climate change. Ethiopia in contrast, has a below average future climate resilience score and very low genomic adaptive capacity score based on the study genotypes collected in the country. These low scores suggest that these genotypes do not show broad climate adaptation and are not resilient to predicted climate change in the country, possibly because the genotypes are narrowly adapted to specific climate conditions (**Figure 2; Figure S4; Figure S5**). This suggests that Ethiopia may need to undertake more significant breeding aimed at adaptation to climate change compared to countries with higher scores.

The indices are capturing different information as seen by the low correlation (**Figure S6**). One advantage of the EGS approach is that it infers likely climate resilience (as opposed to simply making inferences from where a landrace was collected). Additionally, the relationship between phenotype and allelic contributions may be obscured for complex traits, similar to wild tomato (*Lycopersicon pimpinellifolium*) genotypes that can contribute alleles for large fruit size despite their own small-fruited phenotype [34]. This can be extended to environmental adaptation, for example in rice where progeny in a large backcross breeding program often showed submergence and salinity tolerance, among other abiotic stress tolerances, regardless of donor performance [35]. If one relied on collection site data only, one would not be able to detect such mismatches. Ultimately, this information has implications for both national-level strategies in adapting breeding programs and specific breeding program strategy. We outline some of the different decision points when exploring the utility of germplasm collections to be used; these include different decision makers, questions, and results (**Table 1**).

**Table 1.**
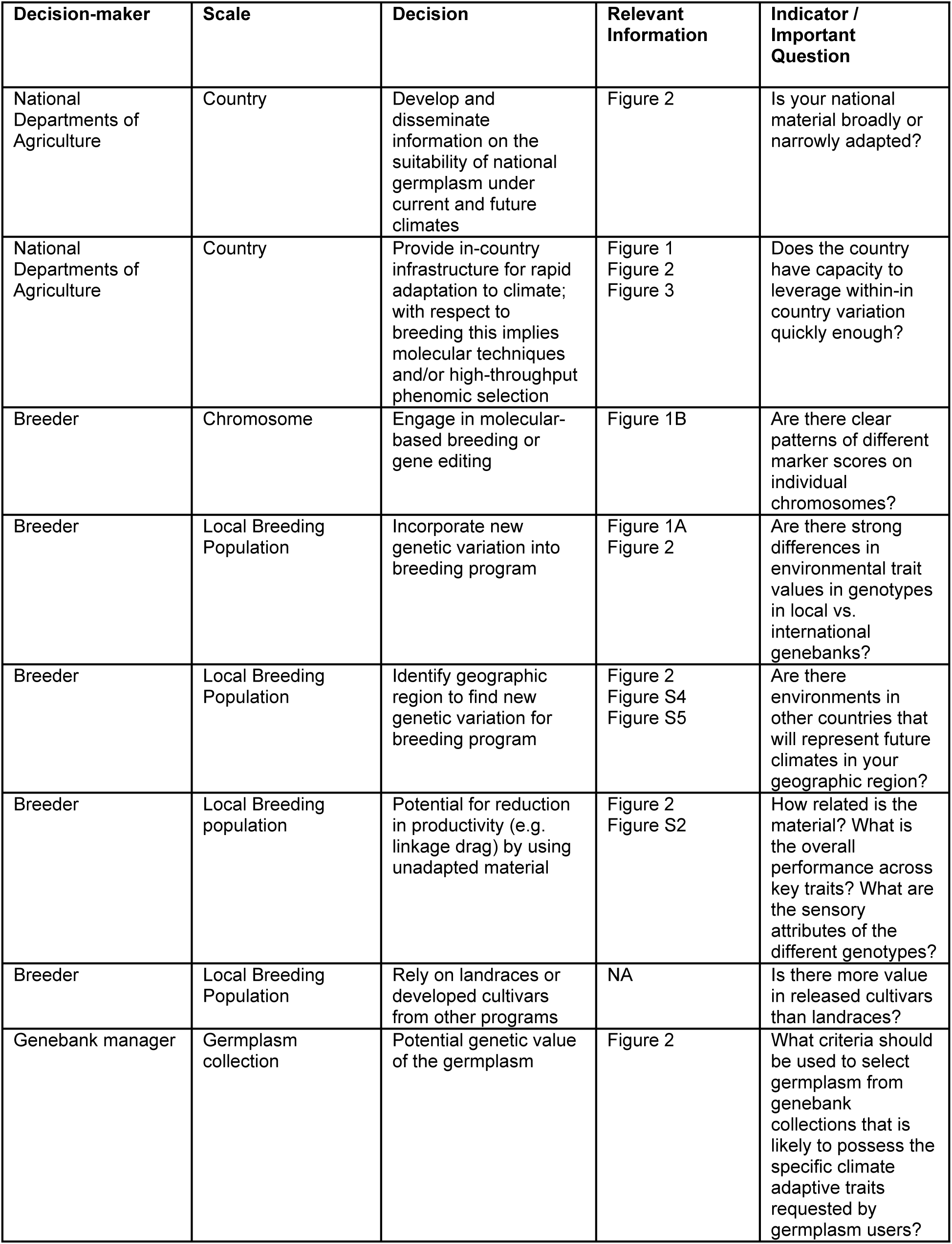
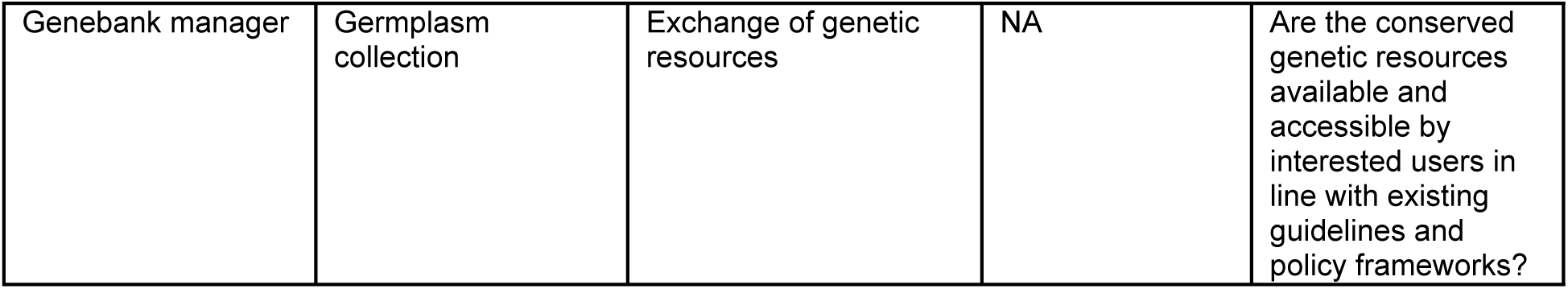
The role of different types of information in decision making for utilizing genetic resources for adaptation to climate change.

| **Decision-maker** | **Scale** | **Decision** | **Relevant Information** | **Indicator / Important Question** |
| --- | --- | --- | --- | --- |
| National Departments of Agriculture | Country | Develop and disseminate information on the suitability of national germplasm under current and future climates | Figure 2 | Is your national material broadly or narrowly adapted? |
| National Departments of Agriculture | Country | Provide in-country infrastructure for rapid adaptation to climate; with respect to breeding this implies molecular techniques and/or high-throughput phenomic selection | Figure 1<br>Figure 2<br>Figure 3 | Does the country have capacity to leverage within-in country variation quickly enough? |
| Breeder | Chromosome | Engage in molecular-based breeding or gene editing | Figure 1B | Are there clear patterns of different marker scores on individual chromosomes? |
| Breeder | Local Breeding Population | Incorporate new genetic variation into breeding program | Figure 1A<br>Figure 2 | Are there strong differences in environmental trait values in genotypes in local vs. international genebanks? |
| Breeder | Local Breeding Population | Identify geographic region to find new genetic variation for breeding program | Figure 2<br>Figure S4<br>Figure S5 | Are there environments in other countries that will represent future climates in your geographic region? |
| Breeder | Local Breeding population | Potential for reduction in productivity (e.g. linkage drag) by using unadapted material | Figure 2<br>Figure S2 | How related is the material? What is the overall performance across key traits? What are the sensory attributes of the different genotypes? |
| Breeder | Local Breeding Population | Rely on landraces or developed cultivars from other programs | NA | Is there more value in released cultivars than landraces? |
| Genebank manager | Germplasm collection | Potential genetic value of the germplasm | Figure 2 | What criteria should be used to select germplasm from genebank collections that is likely to possess the specific climate adaptive traits requested by germplasm users? |
| Genebank manager | Germplasm collection | Exchange of genetic resources | NA | Are the conserved genetic resources available and accessible by interested users in line with existing guidelines and policy frameworks? |

### Identifying climate-adaptive germplasm

Phenotyping germplasm genotypes is logistically difficult in large germplasm collections, especially for complex traits such as tolerance to drought or low nutrient soils. Application of landscape genomic techniques of wild and landrace genotypes to find genes related to climate adaptation provides a way to streamline future phenotypic screening [36-42]. Recently, envirotyping and the integration of environmental data into genomic prediction pipelines has been suggested as a way to exploit climate data [43]. This approach relies on the assumption that landrace genotypes are locally adapted to collection sites due to the long history of selection within a specific geographic region [44]. However, despite the widespread use of these methods to study the genomic basis of local adaptation in wild species and crop landraces there is still limited integration into breeding programs. The present study uses collection information as a proxy for understanding the potential limits of germplasm exchange to adapt to future projected climates. We consider the present scenario, in which the genotypes used in this study are available to requestors internationally, as well as a scenario where germplasm collected in a specific country is only available within said country, for example when international agreements or phytosanitary restrictions restrict germplasm import. Combining FIGS and landscape genomics approaches can create a mechanism to estimate the environmental resilience of any given germplasm panel at any stage of characterization. These metrics provide different complementary ways of understanding resilience and can help with parental choice for a range of traits. Which type of information (genetic or climate projections) will be more useful for breeding programs remains unclear. Bioclimatic data may lead directly to a genotype which could be used for crossing, but using unimproved genotypes could lead to excessive linkage drag. Genetic information permits detection of specific alleles which may already exist in breeding germplasm; however, this requires genotyping of diverse germplasm panels [23]. Further, prediction accuracies could be artificially high because of unaccounted for genetic structure.

In this proof-of-concept study, we have identified specific genotypes with high adaptive values for specific environmental variables, where these genotypes were collected from, and where the accessions are stored (**Table S2**). This approach further permits variation for climate resilience of any germplasm panel to be explored. The approach proposed here could potentially be extended to other crops. However, the extent to which it is applicable depends on the amount of knowledge in that particular species. The techniques explored here require crops with numerous locally adapted landraces, that are grown across a wide range of environments, and for which population structure is weak. For crops that do not meet these criteria (e.g. sunflower, for which the number of suitable landraces is low), then one can potentially make genomic predictions using locally adapted wild genotypes as the training population [39]. The obvious next step is to select parents based on EGS and FIGS and compare which progeny prove superior, which would help provide confidence in this approach.

### The role of germplasm exchange in adapting to climate change

Germplasm exchange is crucial for adapting to the challenges of modern agriculture [45]. The global variation in climate resilience make it clear that responding to climate change will depend on international movement of germplasm to regions where relevant genetic diversity is needed. Sorghum is distributed globally, but the majority of global germplasm is held in two genebanks. The largest collection is held by the ICRISAT in India, while the second largest collection is held in the United States NPGS [46]. While these centers distribute sorghum germplasm all over the world, it is often the case that the countries where these resources were collected do not have substantial collections (**Figure 3**). The current food system, largely based on crops that originated from all over the world with local staples often originating on the other side of the globe, makes modern agriculture highly interdependent [47]. Thus, continued germplasm exchange is a global responsibility to ensure continued food security. Crop improvement for potential future climates requires breeding programs with specific goals based on predicted changes, which vary by geography. To adapt to these diverse changes, breeders need access to the significant genetic variation stored in genebanks globally. Despite facilitated germplasm access through the ITPGRFA, breeders still report that moving germplasm across national borders remains a major limitation in germplasm access [16]. Identifying and characterizing relevant climate adaptation in germplasm collections facilitates the goals of increased global access to these valuable resources.

**Figure 3.**
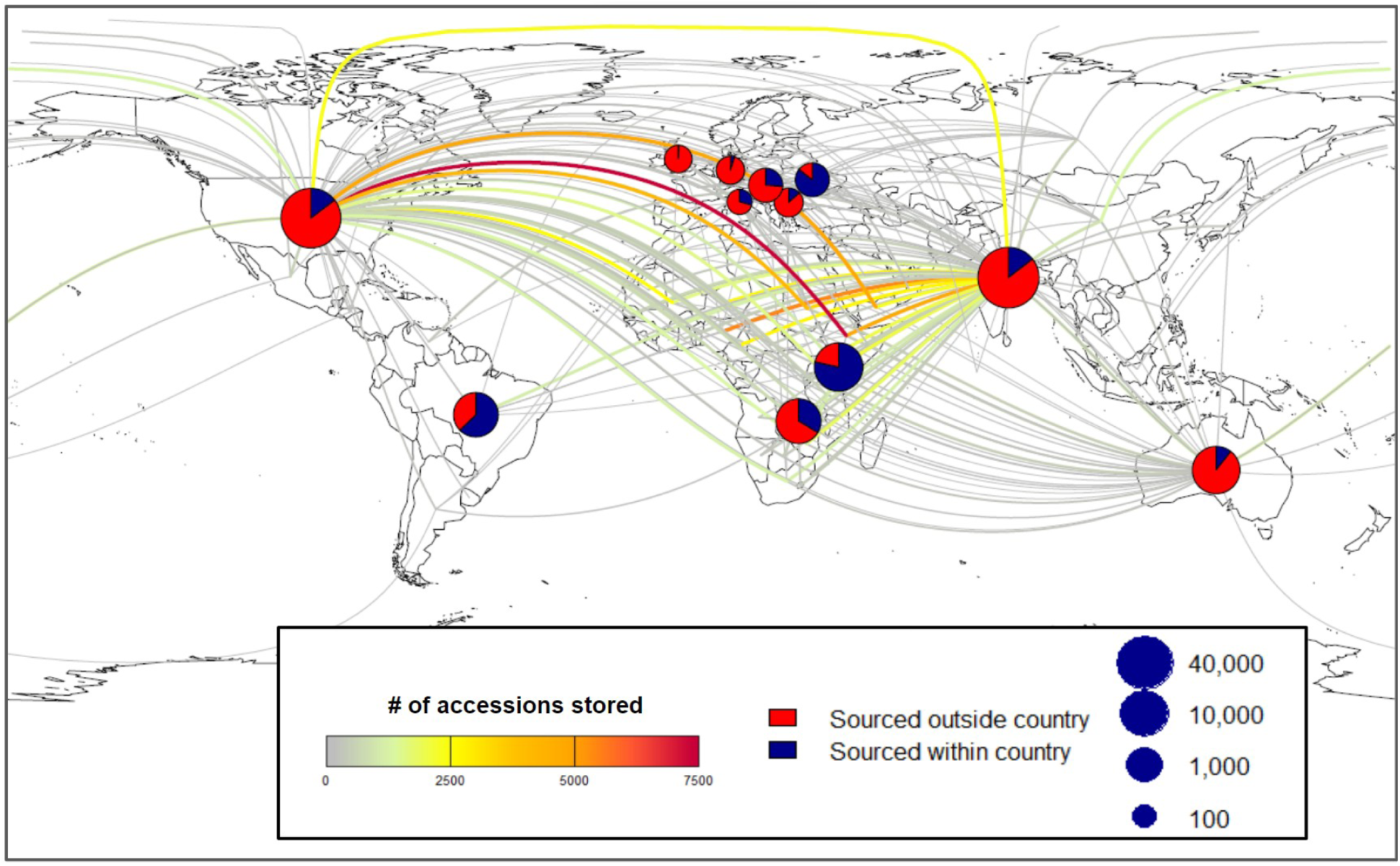
Network map of global sorghum germplasm accessions linking germplasm provenance to repositories where they are currently stored and available for distribution. Pie charts mark repository locations, indicating the percentage of germplasm stored in the repository that have provenance within the country (blue) or outside of the country (red). Color and width of lines represent the number of accessions that are stored from the country of origin within the repository. Data sourced from [genesys-pgr.org].

### Implications for National-level Policy

Through the construction of indices to explore different ways one crop can be adapted across scales, there is a clear pattern of genotypes sourced from different geographic regions having different potential utility (e.g. broad or narrow - **Figure 2**). This has implications for national level food security; some countries with active plant breeding programs will need to have robust mechanisms for germplasm transfer cross national borders, particularly to manage phytosanitary threats such as establishing functional post-entry quarantine infrastructure, increasing the capacity to detect and clean pathogens with the potential to affect local agriculture. For countries lacking breeding infrastructure, open access to germplasm with characterization of climate adaptation will be crucial. This differential national capacity could impact large regional blocks depending on specific scenarios ranging from every-country-for-themselves to completely open. These scenarios are potentially very important to multi-lateral agreements regarding food security. There is also great potential for national governments to embrace the universal right to food and make materials more easily accessible and develop national policies for the implementation of existing and agreed treaties. There is also a need for adequate benefit sharing considering the colonial histories of many conservation organizations [48].

### Conclusion

Climate change is already impacting crop production; taking advantage of past selection and better understanding of climate resilience housed within *ex situ* collections provides actionable strategies for plant breeders and national governments. Germplasm exchange is necessary to adapt crops to climate change. Understanding where needed germplasm resides and what collections need to be made available where will be central to future exchanges. However, better enforcement mechanisms for equity agreements are needed. For this approach to be successful, international systems must be strong and sustainable. The tools outlined in this work can provide an understanding of where climate-adapted germplasm exists, and where it will be most needed under future climates, using publicly available data and accessible methods that can be extended to diverse crop systems.

## Materials and Methods

### Data Acquisition

Genome sequence data from [4] was used for the germplasm panel to conduct environmental Genomic Selection (EGS) on 1,937 sorghum landrace genotypes selected from the ICRISAT (n = 1,093) and NPGS (n = 844) collections. Bioclimatic variables, mean, minimum, and maximum monthly temperature data, and long-term precipitation averages were collected from WorldClim [49]. Soil data were derived from the International Soil Reference and Information Centre (ISRIC) [50-51], with data at six depths for seven aspects of soil (sand content, silt content, clay content, cation exchange capacity, nitrogen, soil organic carbon, bulk density, coarse fragment content). Soil characteristics were aggregated from provided depths (0-5 cm, 5-15 cm, 15-30 cm, 30-60 cm, 60-100 cm, 100-200 cm) into topsoil (0-60 cm) and subsoil (60-200 cm) using weighted means. Topsoil and subsoil categories represent rooting depth for early season and mature flowering stages of sorghum growth.

### Environmental Genomic Selection

Genomic prediction was performed using 404,637 SNPs to predict bioclimatic and biophysical traits (**Table S3-S4**) in order to generate a Genomic Estimated Adaptive Value (GEAV) for each genotype for a given trait (conceptualized as the genetic value for a specific environmental context), in previous work these have been characterized as genomic estimated adaptive values (GEAVs - 32). Four genomic prediction methods were examined: RR-BLUP, G-BLUP with an exponential kernel, G-BLUP with a Gaussian kernel, and BayesCπ. R packages rrBLUP [52] and hibayes [53] were used for model solving (**Table S5)**. Predictions were made for the full study set of 1,937 genotypes using the model established with the summarized data from the 138-genotype mini core collection developed by Upadhyaya et al, 2019 [31] (that were landraces) with georeferenced climate and edaphic variables (**Figure 4**). These 138 georeferenced genotypes were used as the training population. Prediction accuracy was based on Pearson correlation (r(PGE,y)) between the predicted genotypic effects and the observed environmental phenotype. In cross-validation, k-fold validation was performed from k-fold (2, 5, 10) and repeated random sub-sampling (50%, 80%, 90%) scheme to evaluate prediction accuracy. Prediction accuracy was averaged across 50 replications for each combination of genomic prediction method and cross-validation scheme. Environmental traits which were ascribed a prediction accuracy over 50% for the majority of methods examined were used to create a score similar to the future climate resilience score, termed the genomic adaptive capacity score (**Figure S3**). Soil variables showed low prediction accuracy and were excluded from the score calculations. Following assessment of prediction accuracy, RR-BLUP showed the highest prediction accuracies and GEAVs from this model were used for the final score calculations. Within each category (temperature and precipitation), genotype scores were calculated by tallying the number of variables for which the genotype was in the top 5 percent of values for GEAVs, divided by the total number of variables in that category. The two category scores were summed to produce the final genotype score. Country and regional scores were calculated by summing the scores of all genotypes that originated in the given geographical area, divided by the total number of study genotypes originating in the country. Countries with less than 10 genotypes were not included in the country-level score due to unequal sampling distribution. Countries were assigned to regions following the United Nations subregional geoscheme. Chromosomal effects for association with environmental variables were calculated by summing marker effects across each of ten chromosomes for each genotyped sorghum individual in the study. Chromosomal effects were explored through visualizations created in the ComplexHeatMap package in R [54].

**Figure 4.**
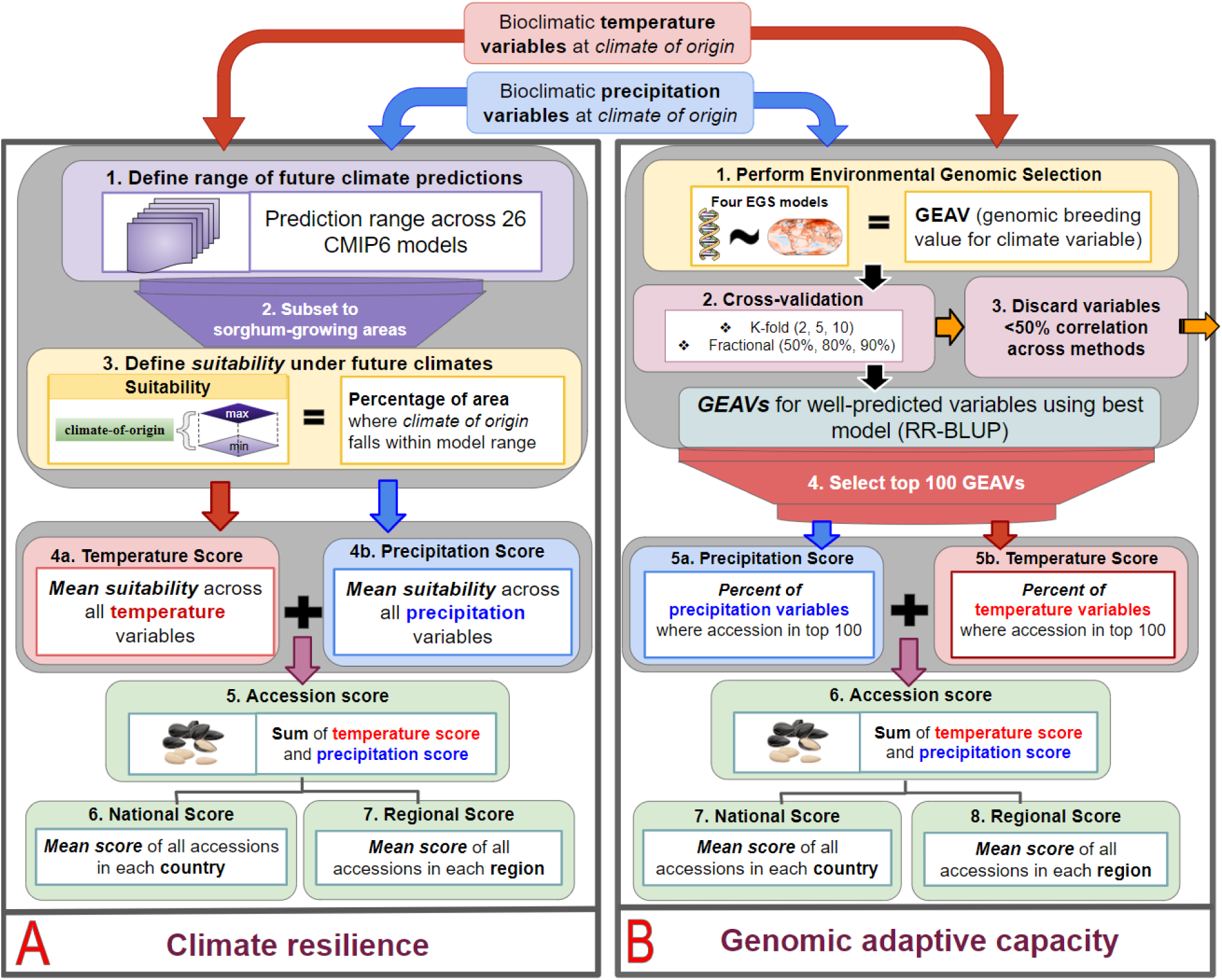
Conceptual figure of how future climate resilience score was calculated. A) Future Climate Score: Resilience score based on CMIP6 future climates models under the SSP 585 scenario using the entire range of values associated with the 23 publicly available Global Circulation Models. Countries with higher scores have more genotypes that originated in the country and are present somewhere in the global germplasm system that have been sourced from environments that are going to be prevalent between 2041-2060. This represents a modified version of the FIGS methodology focusing on identifying the most promising individuals in ex situ collections for pre-breeding. B) Genomic Adaptive Capacity Score: Climate adaptive capacity score based on RR-BLUP values calculated with an environmental genomic selection model using environment-of-origin data as traits associated with 404,627 SNPs scored by Lasky et al, 2016 [4]. Countries with higher scores have a higher proportion of genotypes in the top 100 Genomic Estimated Adaptive Values (GEAVs), or genotypes that are within the top 100 Genomic GEAVs for a larger number of variables. This represents a common technique for selecting candidates for breeding trials.

### Assessment of Core Collection

The ICRISAT sorghum core collection consists of 2,246 sorghum accessions collected globally. More recently, mini core collections have been favored as a way to reduce the number of individuals in a core collection while maintaining genetic diversity. The mini core collection for sorghum was developed by Upadhyaya et al. [31] and consists of 242 accessions, 138 of which are both georeferenced and sequenced. Previously described data was subset and both the future climate resilience score and genomic adaptive capacity score were calculated for the 138 georeferenced accessions. From the full ICRISAT core collection, 1,357 accessions are georeferenced. Environment-of-origin data were collected for these accessions and used to calculate the future climate score and regional and national resilience scores. There were insufficient sequenced core collection genotypes to calculate the genomic adaptive capacity score based on rank of the GEAVs.

### Future Climate Resilience Score (FIGS analog)

The future climate resilience score evaluates the study genotypes for resilience to future climate in their country of origin. The study area encompassed all countries from which genotypes in the study originated, subset to sorghum cropland area from the EarthStat Crop Harvested Area database [55]. A range of future climate scenarios was created for each climatic variable using CMIP6 models under the SSP 585 scenario [56]. Geospatial functions in the terra package and the sf package in R [57-58] were used to find the minimum and maximum value of climatic variables predicted under 23 global climate models for each cell in the study area. Each genotype was evaluated for resilience to future climates for each variable by calculating the percentage of cropland cells within the country of origin where the genotype climate-of-origin values fell between the minimum and maximum values for the future climate predictions [59]. This calculated value was used to calculate two categorical scores, temperature, and precipitation, for an accession by taking the mean of these percentages across 13 temperature variables and 12 precipitation variables. Variable relationships can be seen in **Figure S7**. The overall genotype score was the sum of the two categorical scores. Country scores and regional scores for germplasm climate resilience were found by taking the mean genotype scores across all genotype originating in a given geographical area (**Figure 4**). Data can be found at https://figshare.com/s/da4ad3fe9f749f2135b6 and code can be found at https://github.com/quinncampbell0/sorghum_germplasm_eGS.

## Supporting information

Supplemental able 3

## Supplemental Material

**Figure S1.**
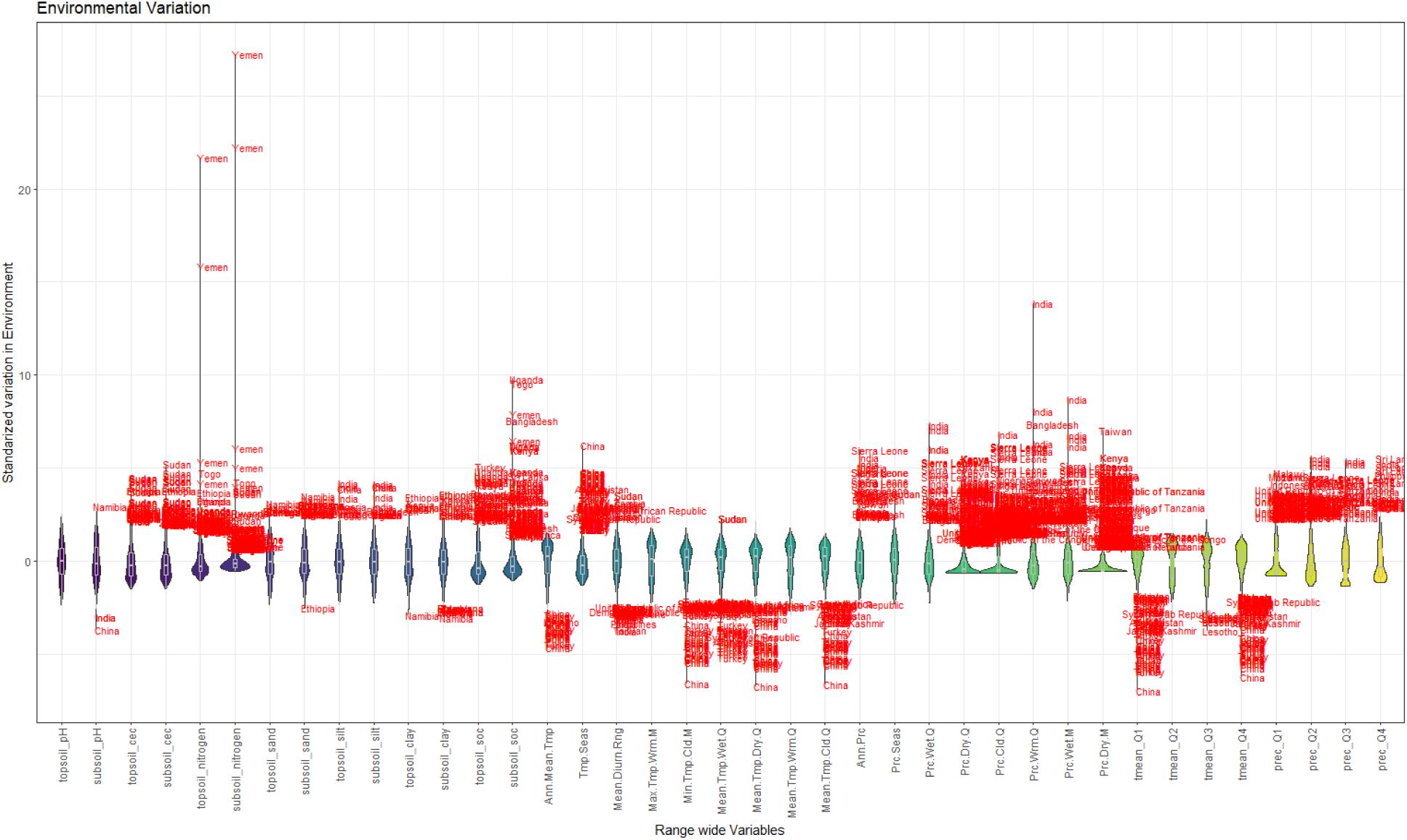
Environmental variation of global collection. Data have been standardized to a mean of 0 and standard deviation of 1.

**Figure S2.**
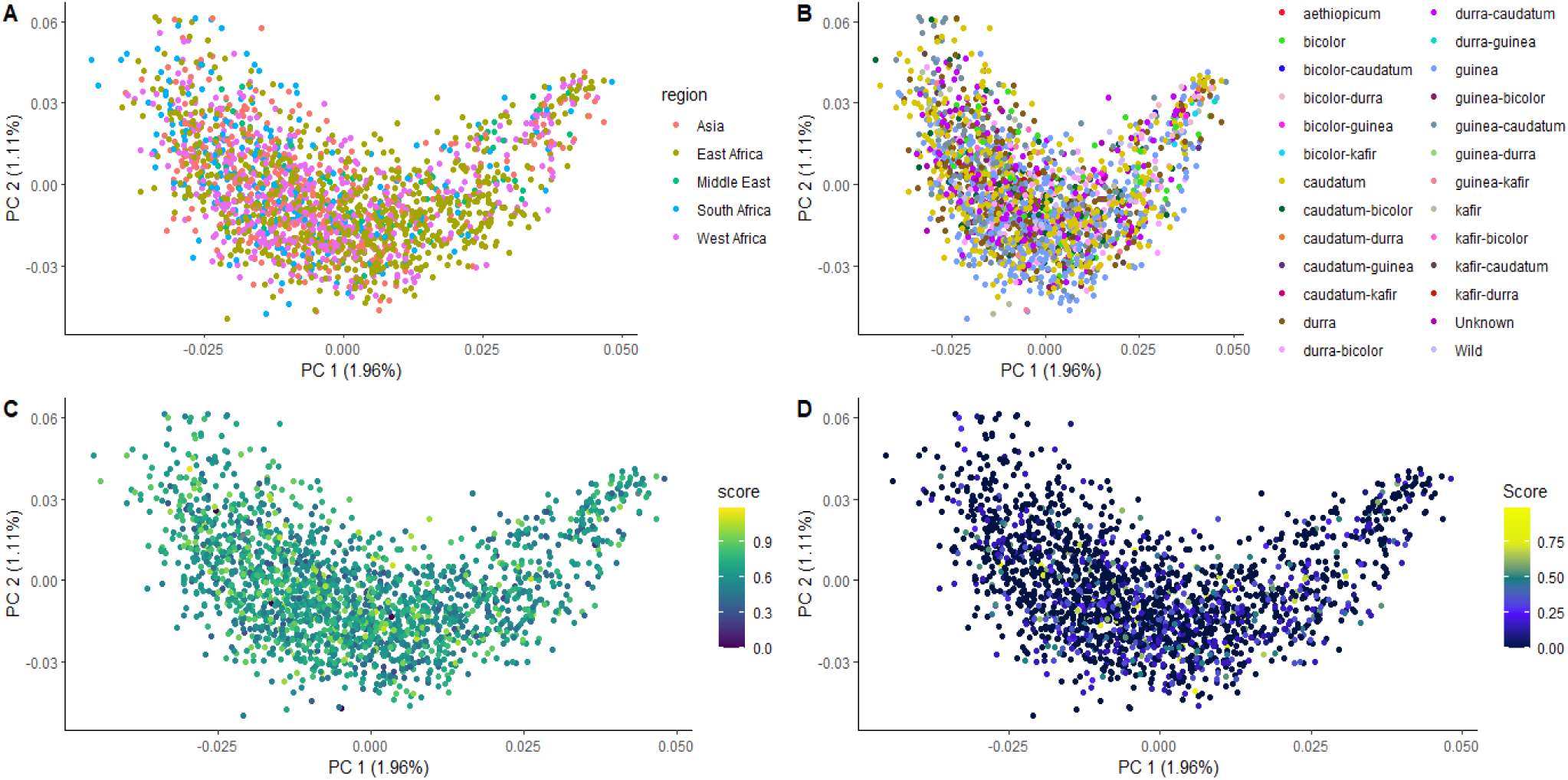
Principal component analysis of SNPs (pruned at 0.2 LD) for global collection with points colored by: A) Geographic region, B) Botanical race, C) Future Climate Resilience Score, and D) Genomic Adaptive Capacity Score

**Figure S3.**
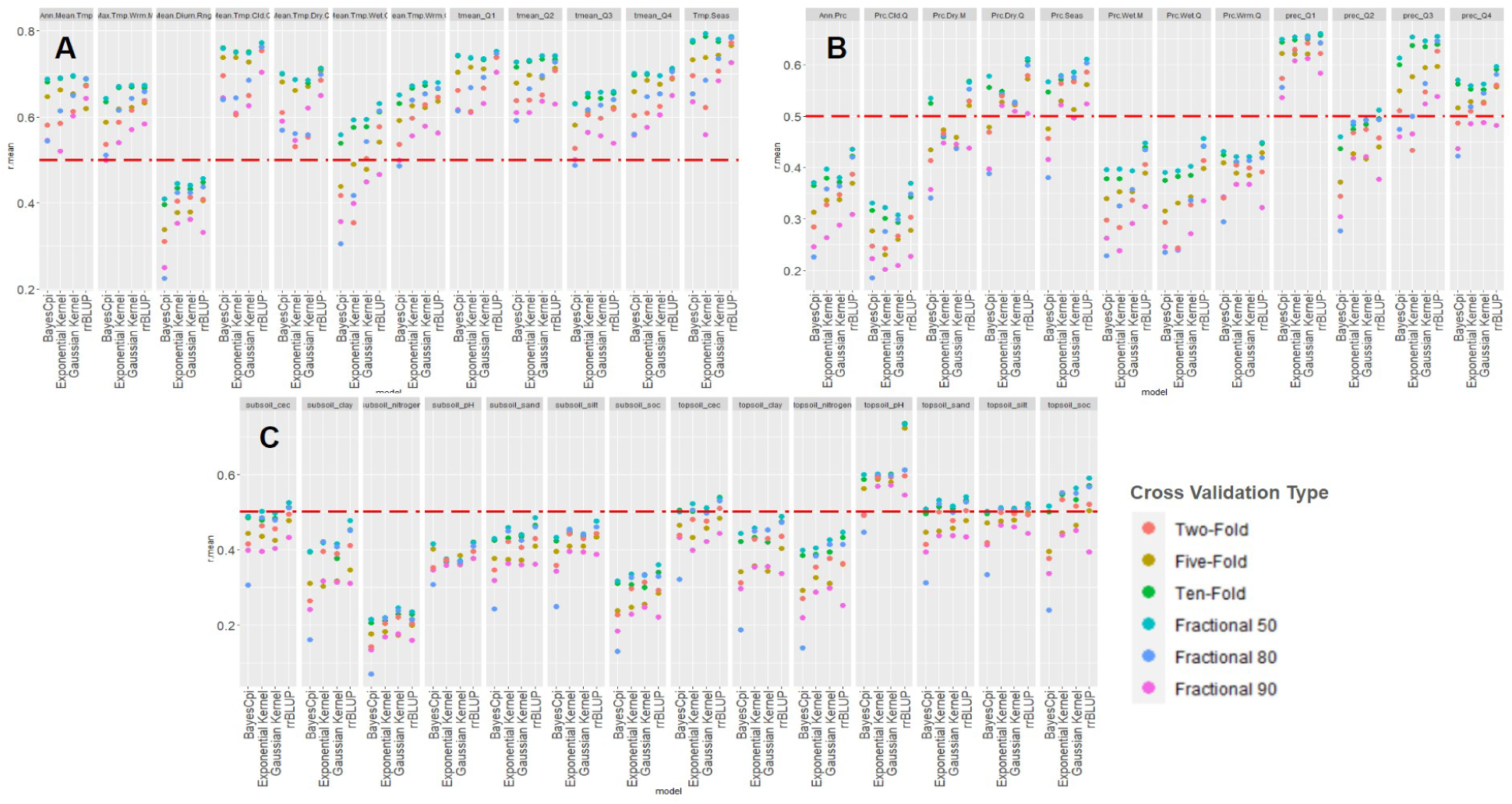
Cross validation of genomic prediction. Four genomic prediction methods (RR-BLUP, G-BLUP with a Gaussian Kernel, G-BLUP with an exponential kernel, and BayesCPi) were evaluated using 6 cross-validation schemes. Training and test sets were subsampled from the mini core collection using k-fold (2, 5, and 10), and repeated random (with 50%, 80%, or 90% as training set) sub-sampling. Prediction accuracy (r(PGE,y)) is reported for all A) temperature, B) precipitation, and C) soil variables.

**Figure S4.**
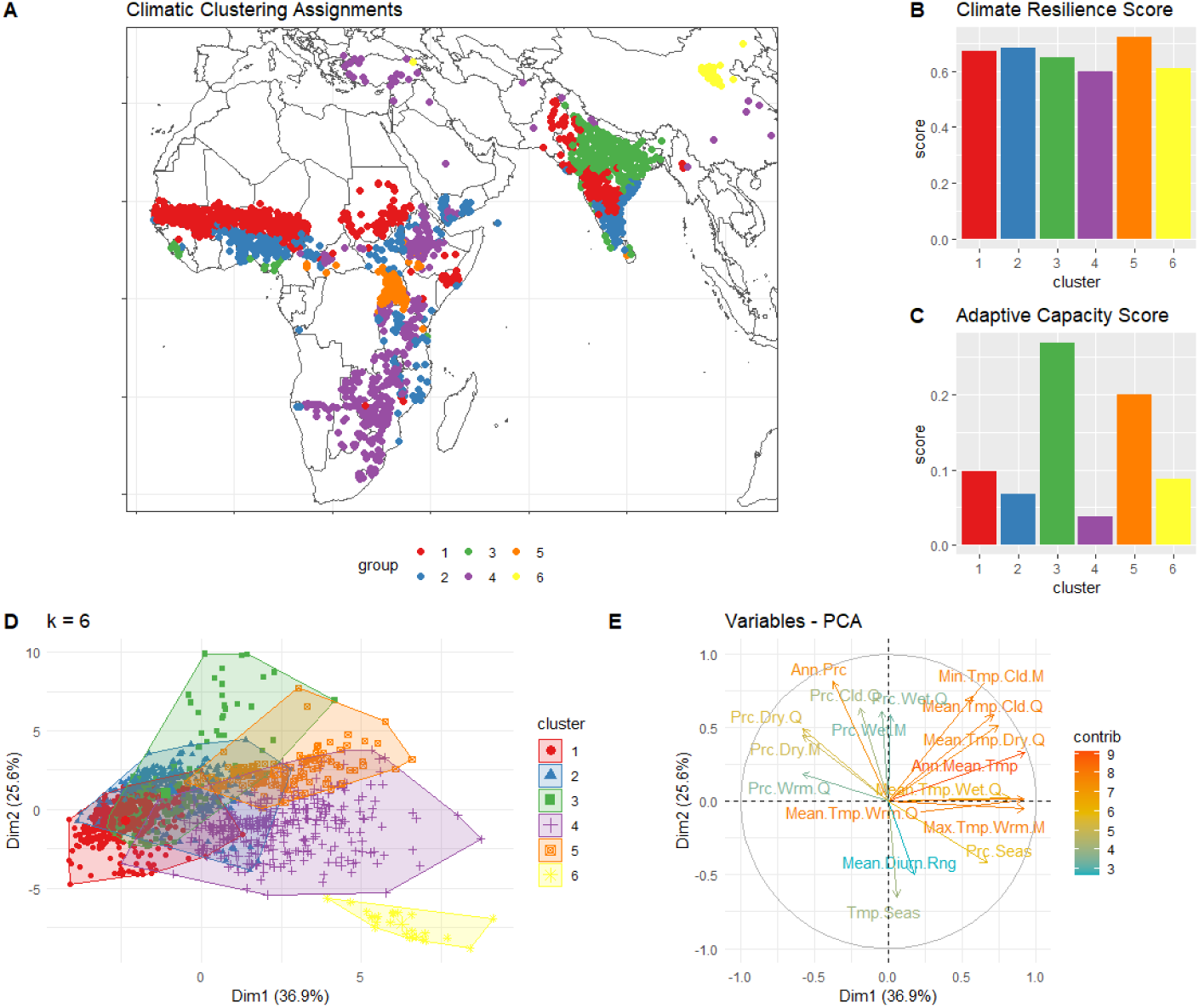
Partitioning score by hierarchical clusters on environment-of-origin data. A) Map of global collection colored by assigned cluster. Bar charts of mean future climate resilience score (B) and mean genomic adaptive capacity score (C) for each climate cluster. Distribution of accessions (D) and variable contribution (E) for the PCA underlying hierarchical clustering assignments. While climate resilience does not vary by cluster, certain climatic clusters show high genomic adaptive capacity scores, possibly because these regions represent extreme environments as quantified by the bioclimatic variables, and these accessions show adaptation to climate extremes.

**Figure S5.**
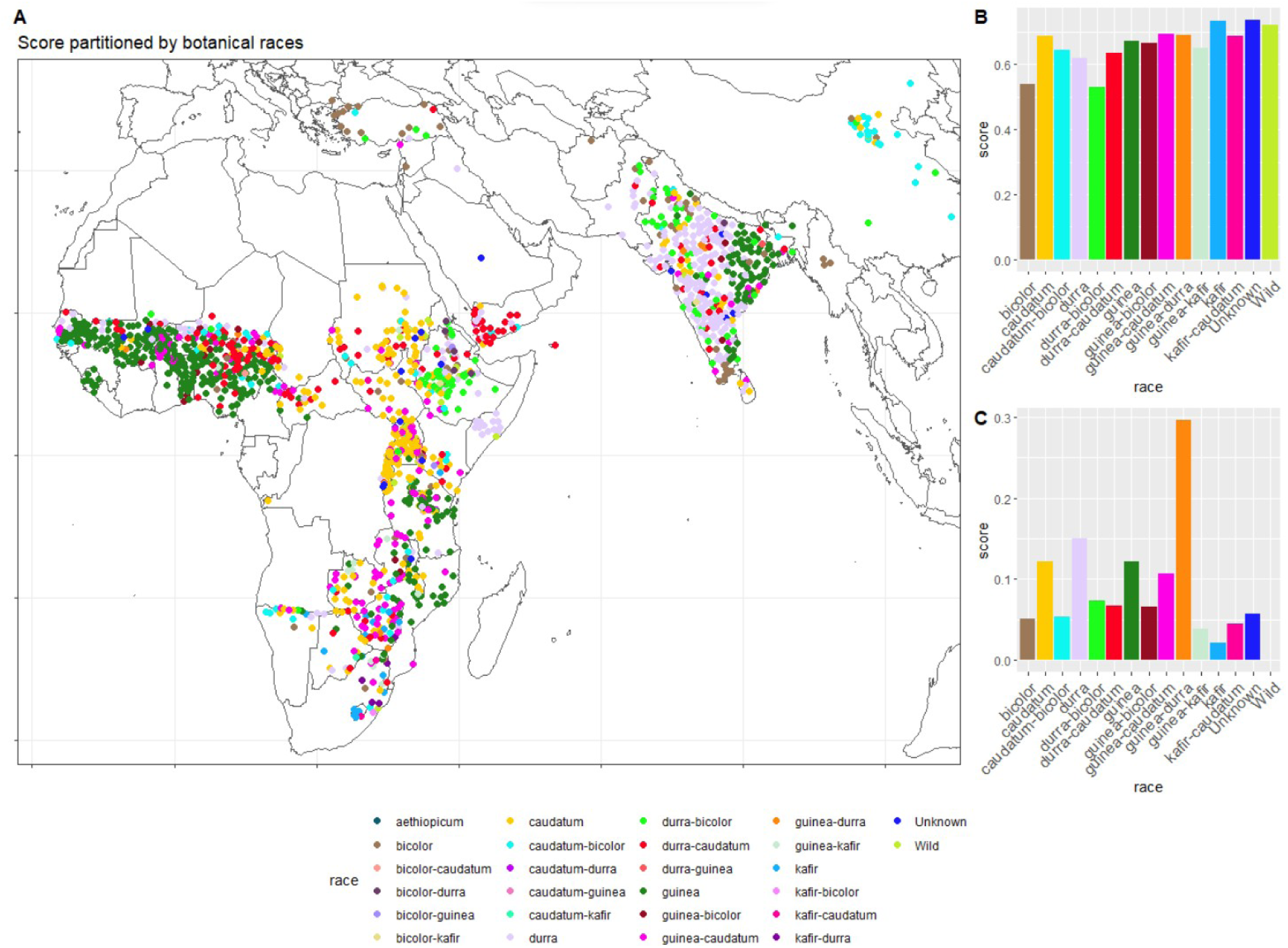
Partitioning of scores by botanical race. Botanical race classifications obtained from Genesys accession passports. Map of accessions colored by botanical race (A) with mean future climate resilience score (B) and mean genomic adaptive capacity score (C) for each botanical race. Little variation is seen across races for climate resilience, likely due to the distribution of these races across geographic areas. However, the guinea-durra race shows very high genomic adaptive capacity, which may be explored for breeding for climate change.

**Figure S6.**
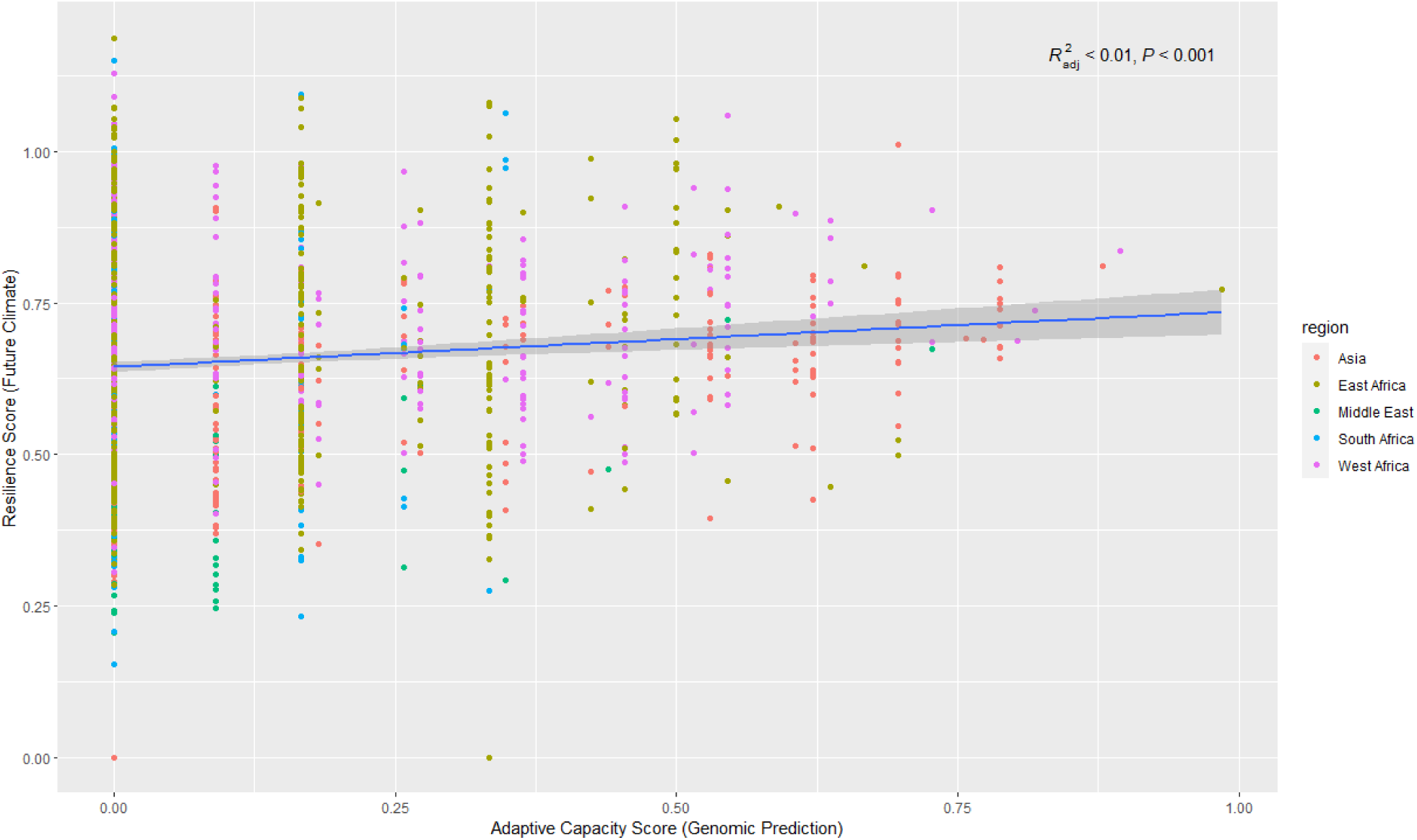
Correlation between the future climate resilience score (FIGS analog) and genomic adaptive capacity score. The lack of a strong relationship suggests that different sources of variation are being captured by each of these indices.

**Figure S7.**
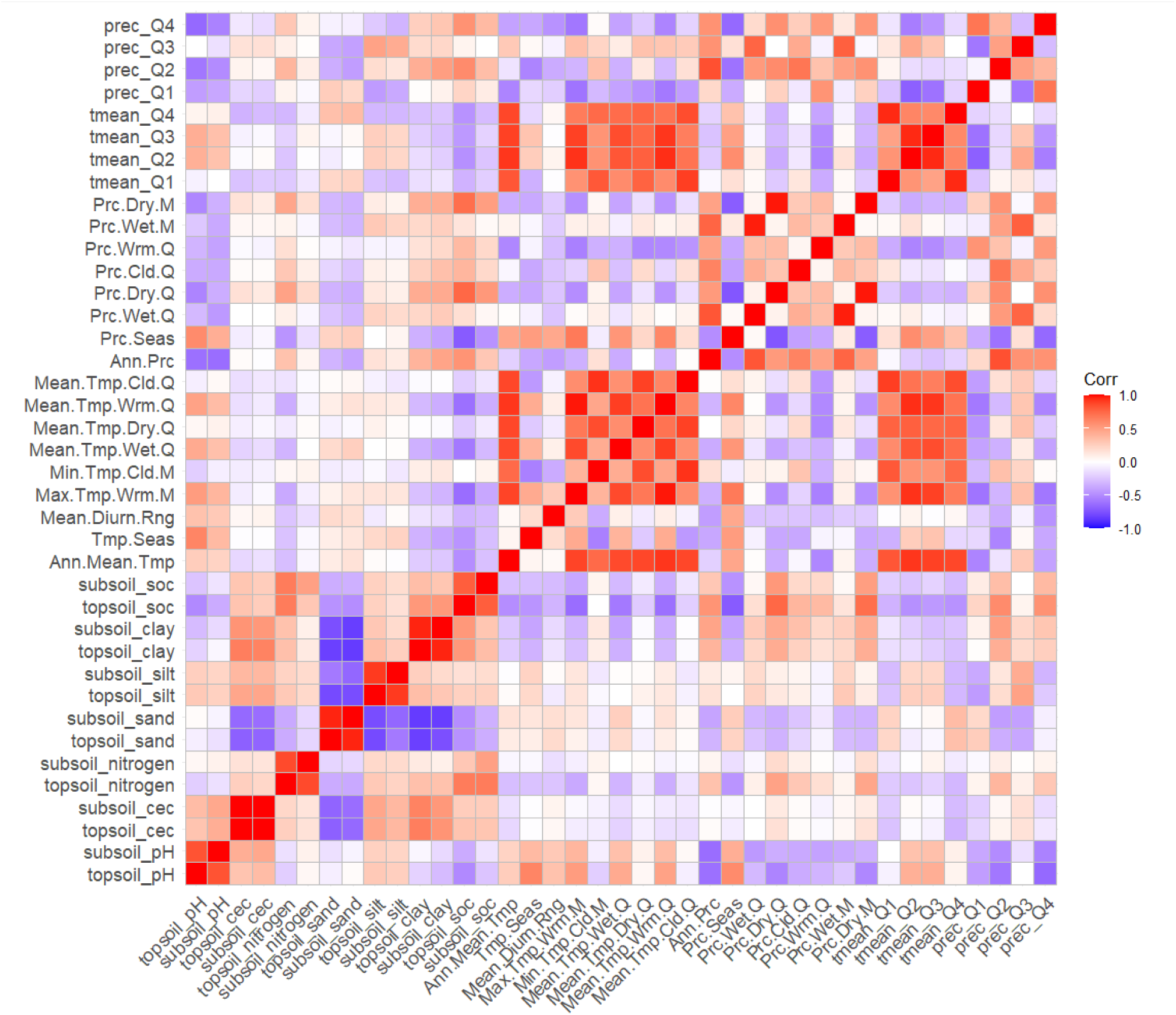
Pairwise correlation between all evaluated climate and edaphic variables used in the study. There are strong relationships between precipitation variables, between temperature variables and between soil properties at different depths.

**Supplemental Table 1.**
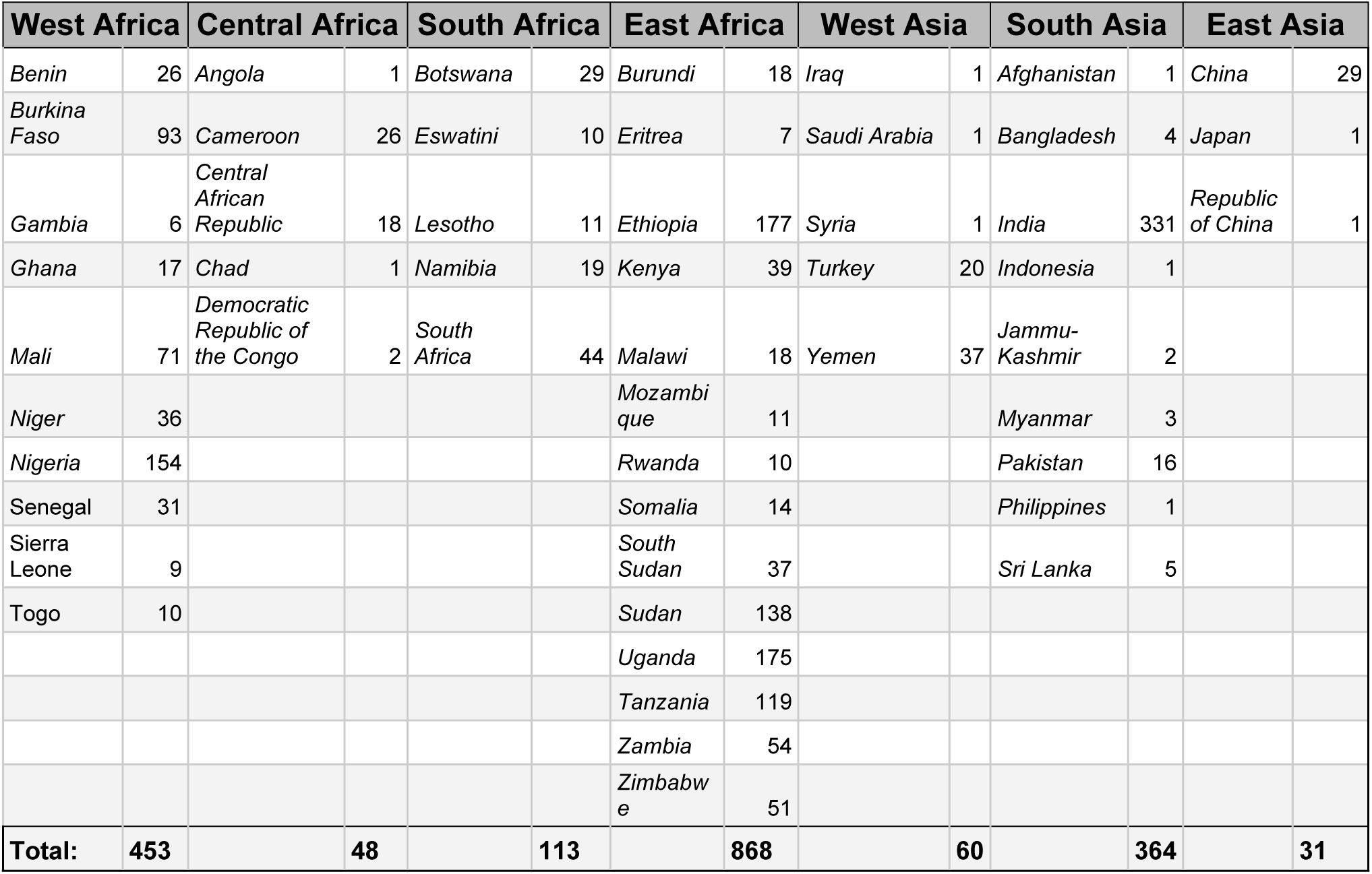
Geographical distribution of the 1,937 accessions in the landrace study collection across countries and regions.

**Supplemental Table 2.** Full table of Genomic Estimated Adaptive Values (GEAVs) from environmental genomic selection using RR-BLUP. Sorting by the environmental variable of interest allows for identifying potential parents with the highest predicted adaptive values for breeding for climatic adaptation.

**Supplemental Table 3.** Geographical data for collecting location of the landrace germplasm collection.

**Supplemental Table 4.** Metadata of environmental variables used in analysis with a full name, units, and category.

**Supplemental Table 5.** Prediction accuracies (mean Pearson’s *r* across 50 runs) for environmental genomic selection of all environmental variables across cross-validation schemes.

## Acknowledgements

We would like to thank Dr. Diane Wang for helpful comments and suggestions on earlier versions of this manuscript. This material is based upon work supported by the National Science Foundation Graduate Research Fellowship Program under Grant Nos. 1842402 and 2236415. Any opinions, findings, and conclusions or recommendations expressed in this material are those of the author(s) and do not necessarily reflect the views of the National Science Foundation.

## Notes

### Competing Interest Statement

The authors have declared no competing interest.

## References

1. Smith, S., Nickson, T. E. & Challender, M. Germplasm exchange is critical to conservation of biodiversity and global food security. Agronomy Journal 113, 2969–2979 (2021).

2. Khoury, C. K., Achicanoy, H. A., Bjorkman, A. D., et al., (2016). Origins of food crops connect countries worldwide. Proceedings of the Royal Society B, 283, 20160792. 10.1098/rspb.2016.0792

3. Khoury, C. K., Brush, S., Costich, D. E., Curry, H. A., de Haan, S., Engels, J. M., et al. (2022). Crop genetic erosion: Understanding and responding to loss of crop diversity. New Phytologist, 233(1), 84–118.

4. Lasky, J. R., Upadhyaya, H. D., Ramu, P., Deshpande, S., Hash, C. T., Bonnette, J., et al. G. P. (2015). Genome-environment associations in sorghum landraces predict adaptive traits. Science advances, 1(6), e1400218.

5. Henry, R. J. (2016). Genomics strategies for germplasm characterization and the development of climate resilient crops. In Crop Breeding (pp. 25-34). Apple Academic Press.

6. Langridge, P., & Waugh, R. (2019). Harnessing the potential of germplasm collections. Nature Genetics, 51(2), 200–201

7. Menamo, T., Kassahun, B., Borrell, A. K., Jordan, D. R., Tao, Y., Hunt, C., & Mace, E. (2021). Genetic diversity of Ethiopian sorghum reveals signatures of climatic adaptation. Theoretical and Applied Genetics, 134, 731–742.

8. Pardey, P. G., Koo, B., Wright, B. D., Van Dusen, M. E., Skovmand, B., & Taba, S. (2001). Costing the conservation of genetic resources: CIMMYT’s ex situ maize and wheat collection. Crop Science, 41(4), 1286–1299.

9. Koo, B., Pardey, P. G., & Wright, B. D. (2003a). The economic costs of conserving genetic resources at the CGIAR centers⋆. Agricultural economics, 29(3), 287–297.

10. Koo, B., Pardey, P., & Wright, B. (2003b). The price of conserving agricultural biodiversity. nature biotechnology,21(2), 126–128.

11. Engels, J. M. M., & Ebert, A. W. (2021). A Critical Review of the Current Global Ex Situ Conservation System for Plant Agrobiodiversity. II. Strengths and Weaknesses of the Current System and Recommendations for Its Improvement. Plants, 10(9), 1904. 10.3390/plants10091904

12. Dempewolf, H., Krishnan, S., & Guarino, L. (2023). Our shared global responsibility: Safeguarding crop diversity for future generations. Proceedings of the National Academy of Sciences, 120(14), e2205768119.10.1073/pnas.2205768119

13. Halewood, M., Chiurugwi, T., Sackville Hamilton, R., Kurtz, B., Marden, E., Welch, E., … & Powell, W. (2018). Plant genetic resources for food and agriculture: opportunities and challenges emerging from the science and information technology revolution. New Phytologist, 217(4), 1407–1419.

14. Kumar, P. L., Cuervo, M., Kreuze, J. F., Muller, G., Kulkarni, G., Kumari, S. G., … & Negawo, A. T. (2021). Phytosanitary interventions for safe global germplasm exchange and the prevention of transboundary pest spread: the role of CGIAR germplasm health units. Plants, 10(2), 328.

15. Ebert, A.W., Engels, J.M., Schafleitner, R., Hintum, T.V. and Mwila, G., 2023. Critical review of the increasing complexity of access and benefit-sharing policies of genetic resources for genebank curators and plant breeders–a public and private sector perspective. Plants, 12(16), p.2992.

16. Galluzzi, G., Seyoum, A., Halewood, M., López Noriega, I., & Welch, E. W. (2020). The Role of Genetic Resources in Breeding for Climate Change: The Case of Public Breeding Programmes in Eighteen Developing Countries. Plants (Basel, Switzerland), 9(9), 1129. 10.3390/plants9091129

17. Deryng, D., Conway, D., Ramankutty, N., Price, J., & Warren, R. (2014). Global crop yield response to extreme heat stress under multiple climate change futures. Environmental Research Letters, 9(3), 034011.

18. Mahaut, L., Pironon, S., Barnagaud, J. Y., Bretagnolle, F., Khoury, C. K., Mehrabi, Z. et al. (2022). Matches and mismatches between the global distribution of major food crops and climate suitability. Proceedings of the Royal Society B, 289(1983), 20221542.

19. Byrne, P. F., Volk, G. M., Gardner, C., Gore, M. A., Simon, P. W., & Smith, S. (2018). Sustaining the future of plant breeding: The critical role of the USDA-ARS National Plant Germplasm System. Crop Science, 58(2), 451–468.

20. Chaloner, T. M., Gurr, S. J., & Bebber, D. P. (2021). Plant pathogen infection risk tracks global crop yields under climate change. Nature Climate Change, 11(8), 710–715.

21. Pironon, S., Etherington, T. R., Borrell, J. S., Kühn, N., Macias-Fauria, M., Ondo, I., … & Willis, K. J. (2019). Potential adaptive strategies for 29 sub-Saharan crops under future climate change. Nature Climate Change, 9(10), 758–763.

22. Brown, A. H. D. (1989). Core collections: a practical approach to genetic resources management. Genome, 31(2), 818–824.

23. Hübner S, Kantar MB. Tapping Diversity From the Wild: From Sampling to Implementation. Front Plant Sci. 2021 Jan 27;12:626565. doi: 10.3389/fpls.2021.626565. PMID: 33584776; PMCID: PMC7873362.

24. Stenberg, J. A., & Ortiz, R. (2021). Focused identification of germplasm strategy (FIGS): polishing a rough diamond. Current Opinion in Insect Science, 45, 1–6.

25. Bragg, J. G., Supple, M. A., Andrew, R. L., & Borevitz, J. O. (2015). Genomic variation across landscapes: insights and applications. New Phytologist, 207(4), 953–967.

26. Lasky, J. R., Josephs, E. B., & Morris, G. P. (2023). Genotype–environment associations to reveal the molecular basis of environmental adaptation. The Plant Cell, 35(1), 125–138.

27. Muleta, K. T., Felderhoff, T., Winans, N., Walstead, R., Charles, J. R., Armstrong, J. S., … & Morris, G. P. (2022). The recent evolutionary rescue of a staple crop depended on over half a century of global germplasm exchange. Science advances, 8(6), eabj4633.

28. Yu, X., Li, X., Guo, T., Zhu, C., Wu, Y., Mitchell, S. E., et al.(2016). Genomic prediction contributing to a promising global strategy to turbocharge gene banks. Nature Plants, 2(10), 1–7.

29. Hickey, L. T., N Hafeez, A., Robinson, H., Jackson, S. A., Leal-Bertioli, S., Tester, M., et al. (2019). Breeding crops to feed 10 billion. Nature biotechnology, 37(7), 744–754.

30. Cudjoe, G. P., Antwi-Agyei, P., & Gyampoh, B. A. (2021). The Effect of Climate Variability on Maize Production in the Ejura-Sekyedumase Municipality, Ghana. Climate, 9(10), Article 10. 10.3390/cli9100145

31. Upadhyaya, H.D., Pundir, R.P.S., Dwivedi, S.L., Gowda, C.L.L., Reddy, V.G. and Singh, S. (2009), Developing a Mini Core Collection of Sorghum for Diversified Utilization of Germplasm. Crop Sci., 49: 1769–1780. 10.2135/cropsci2009.01.0014

32. Cortés, A. J., López-Hernández, F., & Blair, M. W. (2022). Genome–environment associations, an innovative tool for studying heritable evolutionary adaptation in orphan crops and wild relatives. Frontiers in Genetics, 13, 910386.

33. Grenier, C., Hamon, P. and Bramel-Cox, P.J. (2001), Core Collection of Sorghum: II. Comparison of Three Random Sampling Strategies. Crop Science, 41: 241–246. 10.2135/cropsci2001.411241x

34. Lippman, Z., & Tanksley, S. D. (2001). Dissecting the genetic pathway to extreme fruit size in tomato using a cross between the small-fruited wild species Lycopersicon pimpinellifolium and L. esculentum var. Giant Heirloom. Genetics, 158(1), 413–422.

35. Ali, A. J., Xu, J. L., Ismail, A. M., Fu, B. Y., Vijaykumar, C. H. M., Gao, Y. M., Domingo, J., Maghirang, R., Yu, S. B., Gregorio, G., Yanaghihara, S., Cohen, M., Carmen, B., Mackill, D., & Li, Z. K. (2006). Hidden diversity for abiotic and biotic stress tolerances in the primary gene pool of rice revealed by a large backcross breeding program. Field Crops Research, 97(1), 66–76. 10.1016/j.fcr.2005.08.016

36. Bandillo, N. B., Anderson, J. E., Kantar, M. B., Stupar, R. M., Specht, J. E., Graef, G. L., & Lorenz, A. J. (2017). Dissecting the genetic basis of local adaptation in soybean. Scientific reports, 7(1), 17195.

37. Gao, L., Kantar, M. B., Moxley, D., Ortiz-Barrientos, D., & Rieseberg, L. H. (2023). Crop Adaptation to Climate Change: An Evolutionary Perspective. Molecular Plant.

38. Lei, L., Poets, A. M., Liu, C., Wyant, S. R., Hoffman, P. J., Carter, C. K., et al., (2019). Environmental association identifies candidates for tolerance to low temperature and drought. G3: Genes, Genomes, Genetics, 9(10), 3423-3438.

39. Neyhart, J. L., Kantar, M. B., Zalapa, J., & Vorsa, N. (2022). Genomic-environmental associations in wild cranberry (Vaccinium macrocarpon Ait.). G3, 12(10), jkac203.

40. Todesco, M., Owens, G. L., Bercovich, N., Légaré, J. S., Soudi, S., Burge, D. O., … & Rieseberg, L. H. (2020). Massive haplotypes underlie ecotypic differentiation in sunflowers. Nature, 584(7822), 602–607.

41. Wang, D. R., Kantar, M. B., Murugaiyan, V., & Neyhart, J. (2023). Where the wild things are: Genetic associations of environmental adaptation in the Oryza rufipogon species complex. G3: Genes, Genomes, Genetics, jkad128.

42. Pyhäjärvi, T., Hufford, M. B., Mezmouk, S., & Ross-Ibarra, J. (2013). Complex patterns of local adaptation in teosinte. Genome biology and evolution, 5(9), 1594–1609.

43. Costa-Neto, G., Galli, G., Carvalho, H. F., Crossa, J., & Fritsche-Neto, R. (2021). EnvRtype: a software to interplay enviromics and quantitative genomics in agriculture. G3, 11(4), jkab040.

44. Villa, T. C. C., Maxted, N., Scholten, M., & Ford-Lloyd, B. (2005). Defining and identifying crop landraces. Plant genetic resources, 3(3), 373–384.

45. Laird, S., Wynberg, R., Rourke, M., Humphries, F., Muller, M. R., & Lawson, C. (2020). Rethink the expansion of access and benefit sharing. Science, 367(6483), 1200–1202.

46. Bramel P., Kresovich S., and Giovannini P. 2022. Global Strategy for the Conservation and Use of Sorghum (Sorghum bicolor (L.) Moench) Genetic Resources. Global Crop Diversity Trust. Bonn, Germany. DOI: 10.5281/zenodo.8192869

47. Khoury, C. K., Achicanoy, H. A., Bjorkman, A. D., Navarro-Racines, C., Guarino, L., Flores-Palacios, X., Engels, J. M. M., Wiersema, J. H., Dempewolf, H., Sotelo, S., Ramírez-Villegas, J., Castañeda-Álvarez, N. P., Fowler, C., Jarvis, A., Rieseberg, L. H., & Struik, P. C. (2016). Origins of food crops connect countries worldwide. Proceedings of the Royal Society B: Biological Sciences, 283(1832), 20160792. 10.1098/rspb.2016.0792

48. Lenzner, B., Latombe, G., Schertler, A., Seebens, H., Yang, Q., Winter, M., et al. (2022). Naturalized alien floras still carry the legacy of European colonialism. Nature Ecology & Evolution, 6(11), 1723–1732.

49. Fick, S. E., & Hijmans, R. J. (2017). WorldClim 2: new 1-km spatial resolution climate surfaces for global land areas. International journal of climatology, 37(12), 4302–4315.

50. Hengl, T., Mendes de Jesus, J., Heuvelink, G. B., Ruiperez Gonzalez, M., Kilibarda, M., Blagotić, A., et al. (2017). SoilGrids250m: Global gridded soil information based on machine learning. PLoS one, 12(2), e0169748.

51. Hengl, T., de Jesus, J. M., MacMillan, R. A., Batjes, N. H., Heuvelink, G. B., Ribeiro, E., et al. (2014). SoilGrids1km—global soil information based on automated mapping. PloS one, 9(8), e105992.

52. Endelman, J.B. 2011. Ridge regression and other kernels for genomic selection with R package rrBLUP. Plant Genome 4:250–255.

53. Yin L, Zhang H, Liu X (2022). hibayes: Individual-Level, Summary-Level and Single-Step Bayesian Regression Model. R package version 2.0.0, <https://CRAN.R-project.org/package=hibayes>.

54. Gu, Z. (2016) Complex heatmaps reveal patterns and correlations in multidimensional genomic data. Bioinformatics. DOI: 10.1093/bioinformatics/btw313.

55. Monfreda, C., Ramankutty, N., and Foley, J. A. (2008). Farming the planet: 2. Geographic distribution of crop areas, yields, physiological types, and net primary production in the year 2000. Global Biogeochem. Cycles. 22. GB1022. doi:10.1029/2007GB002947.

56. Stockhause, M., Matthews, R., Pirani, A., Treguier, A. M., & Yelekci, O. (2021, April). CMIP6 data documentation and citation in IPCC’s Sixth Assessment Report (AR6). In EGU General Assembly Conference Abstracts (pp. EGU21-2886).

57. Hijmans R (2022). _terra: Spatial Data Analysis_. R package version 1.6–47, <https://CRAN.R-project.org/package=terra>.

58. Pebesma, E., 2018. Simple Features for R: Standardized Support for Spatial Vector Data. The R Journal 10 (1), 439–446, 10.32614/RJ-2018-009

59. Potapov, P., Turubanova, S., Hansen, M. C., Tyukavina, A., Zalles, V., Khan, A., Song, X.-P., Pickens, A., Shen, Q., & Cortez, J. (2022). Global maps of cropland extent and change show accelerated cropland expansion in the twenty-first century. Nature Food, 3(1), Article 1. 10.1038/s43016-021-00429-z

