## supplemental Table 1 for "Climate resilience conserved in global germplasm repositories: Picking the most promising parents for agile plant breeding"

**Supplemental Table 1.** *Geographical distribution of the 1,937 accessions in the landrace study collection across countries and regions.*

| **West Africa** | | **Central Africa** | | **South Africa** | | **East Africa** | | **West Asia** | | **South Asia** | | **East Asia** | |
| --- | --- | --- | --- | --- | --- | --- | --- | --- | --- | --- | --- | --- | --- |
| *Benin* | 26 | *Angola* | 1 | *Botswana* | 29 | *Burundi* | 18 | *Iraq* | 1 | *Afghanistan* | 1 | *China* | 29 |
| *Burkina Faso* | 93 | *Cameroon* | 26 | *Eswatini* | 10 | *Eritrea* | 7 | *Saudi Arabia* | 1 | *Bangladesh* | 4 | *Japan* | 1 |
| *Gambia* | 6 | *Central African Republic* | 18 | *Lesotho* | 11 | *Ethiopia* | 177 | *Syria* | 1 | *India* | 331 | *Republic of China* | 1 |
| *Ghana* | 17 | *Chad* | 1 | *Namibia* | 19 | *Kenya* | 39 | *Turkey* | 20 | *Indonesia* | 1 |  |  |
| *Mali* | 71 | *Democratic Republic of the Congo* | 2 | *South Africa* | 44 | *Malawi* | 18 | *Yemen* | 37 | *Jammu-Kashmir* | 2 |  |  |
| *Niger* | 36 |  |  |  |  | *Mozambique* | 11 |  |  | *Myanmar* | 3 |  |  |
| *Nigeria* | 154 |  |  |  |  | *Rwanda* | 10 |  |  | *Pakistan* | 16 |  |  |
| Senegal | 31 |  |  |  |  | *Somalia* | 14 |  |  | *Philippines* | 1 |  |  |
| Sierra Leone | 9 |  |  |  |  | *South*  *Sudan* | 37 |  |  | *Sri Lanka* | 5 |  |  |
| Togo | 10 |  |  |  |  | *Sudan* | 138 |  |  |  |  |  |  |
|  |  |  |  |  |  | *Uganda* | 175 |  |  |  |  |  |  |
|  |  |  |  |  |  | *Tanzania* | 119 |  |  |  |  |  |  |
|  |  |  |  |  |  | *Zambia* | 54 |  |  |  |  |  |  |
|  |  |  |  |  |  | *Zimbabwe* | 51 |  |  |  |  |  |  |
| **Total:** | **453** |  | **48** |  | **113** |  | **868** |  | **60** |  | **364** |  | **31** |
